# Foundation Model Enables Interpretable Open and Error-Tolerant Searching for Mass Spectrometry-Based Proteomics

**DOI:** 10.1101/2021.12.01.470818

**Authors:** Tom Altenburg, Thilo Muth, Patrick van Zalm, Hanno Steen, Bernhard Y. Renard

## Abstract

Mass spectrometry-based proteomics allows to study all proteins of a sample on a molecular level. However, mass spectra are noisy and contain complex patterns, making them inherently challenging to analyze with purely algorithmic approaches. In terms of the protein sequence landscape, most recent bottom-up MS-based proteomics studies consider either a diverse pool of post-translational modifications, employ large databases – as in metaproteomics or proteogenomics, study multiple isoforms of proteins, include unspecific cleavage sites or even combinations thereof. All this makes peptide and protein identifications challenging due to sheer size of the search space. To cope with this two-sided challenge, i.e. the complexity of real spectra and the search space size, we present a foundation model, called yHydra, that jointly embeds spectra and peptides. This allows us to implement various downstream tasks and search modes in Euclidean space. In particular, we implement an open search which allows to query multiple ten-thousands of spectra against millions of peptides. Furthermore, we implement an error-tolerant search for identifying additional proteoforms that are not included in off-the-shelf reference proteomes. Our foundation model provides meaningful embeddings, as we interpret learned peptide embeddings in comparison to the peptide’s physico-chemical properties. yHydra’s open search, assigns delta masses to each identification which allows to unrestrictedly characterize post-translational modifications. The error-tolerant mode of yHydra can be used as post-processing to existing search engines or as a standalone. yHydra is evaluated on several real life data sets for the identification of modified protein sequences and shows up to 25% increase in protein identification at constant false discovery rate compared to the current state-of-the-art.

**Availability:** (under MIT license) https://gitlab.com/dacs-hpi/yHydra

## Introduction

Peptide sequences and their corresponding fragmentation mass spectra have been the two pivotal peptide representations in mass spectrometry (MS)-based proteomics up to date [1]. A single MS experiment can produce up to one terabyte of data, mostly peptide information represented as tandem mass spectra when using recent instrumentation. To identify actual peptide sequences from spectra a conventional proteomics search engine is used in most cases [2]. Such a search engine constructs theoretical spectra derived from a context-matched reference proteome to identify the actual peptide sequence by performing comparisons in spectrum space. Each comparison is effortful as it involves the construction of spectra, peak matching, intensity summation, tie-breaking, score calculation and post-processing. The construction of spectra itself is challenging as real spectra have missing peaks, noise peaks (i.e. other than peptide-originating peaks), depict complex patterns due to the overlay of isotopic distributions of fragments, and peak heights are non-trivially characteristic to the underlying peptide. Here, we present a foundation model, called yHydra, that jointly embeds peptides and tandem mass spectra as vectors into the same Euclidean space. This way comparisons can be performed efficiently using Euclidean (L2-)distances between spectrum embeddings and peptide embeddings. yHydra is a foundation model and enables us to tackle various downstream tasks and implement multiple search modes. Specifically, we implement three search modes including closed, open and error-tolerant search modes.

Recently, various machine learning-based approaches have been introduced to improve the downstream data analysis in MS-based proteomics [3, 4]. Most of these approaches improve specific aspects of the aforementioned conventional database search. For example, fragment intensity prediction substantially improves search outcomes by using recent deep learning techniques [5, 6]. Post-processing, so-called rescoring, has been very successful by using semi-supervised learning [7, 8]. Auxiliary peptide properties such as ion mobility or retention time, acquired during the MS-experiment, can be predicted from deep learning models to further improve rescoring [9, 10]. Still, all these methods operate within the boundaries of a conventional database search and are conceptually limited by operating in spectrum space.

A major challenge in MS-based proteomics are proteoforms. These protein variants arise from mutations, splicing, processing, and/or post-translational modifications (PTMs). All these deviations from the reported primary sequence are relevant for physiological processes but also involved in human diseases [11]. Taking proteoforms into account means searching against more complex and larger search spaces. This can be achieved via an open search where a considerably wide mass window allows for a more flexible and larger search context per spectrum. A collection of algorithmic approaches for open searches exist, including index– [12, 13], tag– [14, 15] or spectral library-based methods [16]. All these approaches are purely algorithmic. In contrast, yHydra as a foundation model was trained on real spectra, providing a more nuanced spectrum representation, which results in improved peptide identification. The above mentioned algorithms fall short as they only reflect limited feature sets from manual engineering. Our approach, allows to study proteoforms from three angles. First, by implementing an open search capturing the heuristic nature of real spectra through the embeddings. Second, by constructing a delta mass profile, which summarizes differences between proteoform masses and canonical peptide masses, indicative of PTMs or mutations. Third, by investigating particular regions of the embedding space for potential enrichment of PTMs.

There exist deep learning approaches for specific tasks in MS-based proteomics, including the detection of modified peptides [17], spectrum-to-peptide translation (so-called *de novo* sequencing) [18, 19, 20, 21, 22] and a pairwise spectrum embedding for clustering [23]. There exists a cross-modal embedding for learning a similarity metric between tandem mass spectra and peptides [24]. Such learned similarity metric allows heuristic scoring of peptide spectrum matches (PSMs) but is limited to closed searches only.

Altogether, all the above methods are dedicated but limited to single tasks. None of these methods aims to provide a universal basis of how to compare spectra and peptides in a unified and interpretable manner. To mitigate these shortcomings, we provide a foundation model via learned joint embeddings of spectra and peptides. Having a unified view on both representations allows to implement various search modes and types of downstream analyses in embeddings space, of which we implemented three search modes and additionally fine-tuned our model to cope with the two most commonly used instruments in MS-based proteomics and also cover tryptic and non-tryptic peptides.

Here, we present our foundation model yHydra, consisting of a Spectrum Transformer and a Peptide Transformer. The Transformer model architecture has been successful in various domains due to its generalization capabilities [25]. Our approach is inspired by one of the first true foundation models, called CLIP, originally embedding images and text [26]. For training, we used 67 different proteomics repositories containing nearly 20 million identified peptide spectrum matches (PSMs). We show that the high resolution of modern mass spectrometers is well covered by our peak encodings, allowing to capture all information of real spectra. Next, we verify our approach by interpreting our embeddings using Uniform Manifold Approximation and Projection (UMAP). This demonstrates that learned embeddings are meaningful, as the learned manifold reflects physico-chemical properties of the underlying peptides. Once trained, we use the foundation model to implement specific sub-tasks. We start by implementing a closed and open search using a GPU-accelerated k-NN search library as backend [27]. Using our open search we were able to build a delta mass profile for characterizing PTMs in the sample. As an additional sub-task, we implement an error-tolerant search mode by implementing a dedicated beam search in our embedding space. Visualizing the reward matrix based on the gradient towards the decoded peptide sequence during beam search allowed further model interpretability. Having an error-tolerant mode enabled us to identify new proteoforms due to genetic variations. In particular, we searched samples of a monoclonal antibody and chimpanzee plasma.

Protein sequencing of monoclonal antibodies is challenging because of their variable regions [28]. These regions result from immune adaptations and are unique per individuum. Here, we use our error-tolerant search to identify peptides of a tryptic digest of a monoclonal antibody and evaluate our approach by searching against publicly available antibody sequences while comparing peptide identifications to the actual fully assembled antibody sequence as a ground truth.

Proteomes of many non-human primates, such as chimpanzee (*Pan troglodytes*), are not yet fully annotated. A better understanding of their proteomes can aid drug development and research. However, their genomes have been sequenced and corresponding proteins have been suggested [29, 30]. Here, we perform a cross-species error-tolerant search of tryptic peptides from a chimpanzee plasma sample. Therefore, we evaluate our approach by searching against the canonical human proteome and comparing to the genome-derived isoforms of the predicted chimpanzee proteome. Thereby, we capture potential proteoforms due to genetic variations between the proteomes of both species.

## Results

### yHydra learns to jointly embed peptides and MS/MS spectra

To gain a foundation model we trained a deep learning model that jointly embeds peptides and fragmentation spectra. In particular, yHydra embeds both representations into a joint embedding space of real-valued vectors in an Euclidean space (Fig. 1A). Having such a metric between fixed-sized vectors allows computationally inexpensive comparisons between embeddings. To gain embeddings, we jointly trained dedicated Transformer models (one for each of the two domains) by providing a large collection of previously identified peptide spectrum matches (see Methods section). During training, yHydra learns an embedding such that the Euclidean distance between both embeddings of a PSM is small compared to the distances of any mismatched paired embeddings (i.e. by swapping peptides and spectra of PSMs within each mini-batch). Or in other words, to find embeddings such that the diagonal of the pairwise Euclidean distance matrix (Fig. 1A, bottom right) is minimized while the off-diagonal of this matrix is maximized. Note, after both embedders (respectively, Spectrum Transformer and Peptide Transformer) have been trained they can be used separately and independently while both embed into the same joint space. In particular, we implement an open search by embedding MS/MS spectra to search them against embeddings of digested peptides from a proteome database (Fig. 1B). Because both embeddings are real-valued vectors of the same fixed size that live in the same joint embedding space we can make use of highly optimized algorithms, such as the GPU-accelerated k-nearest neighbor (k-NN) search [27], as we demonstrate below.

**Figure 1:**
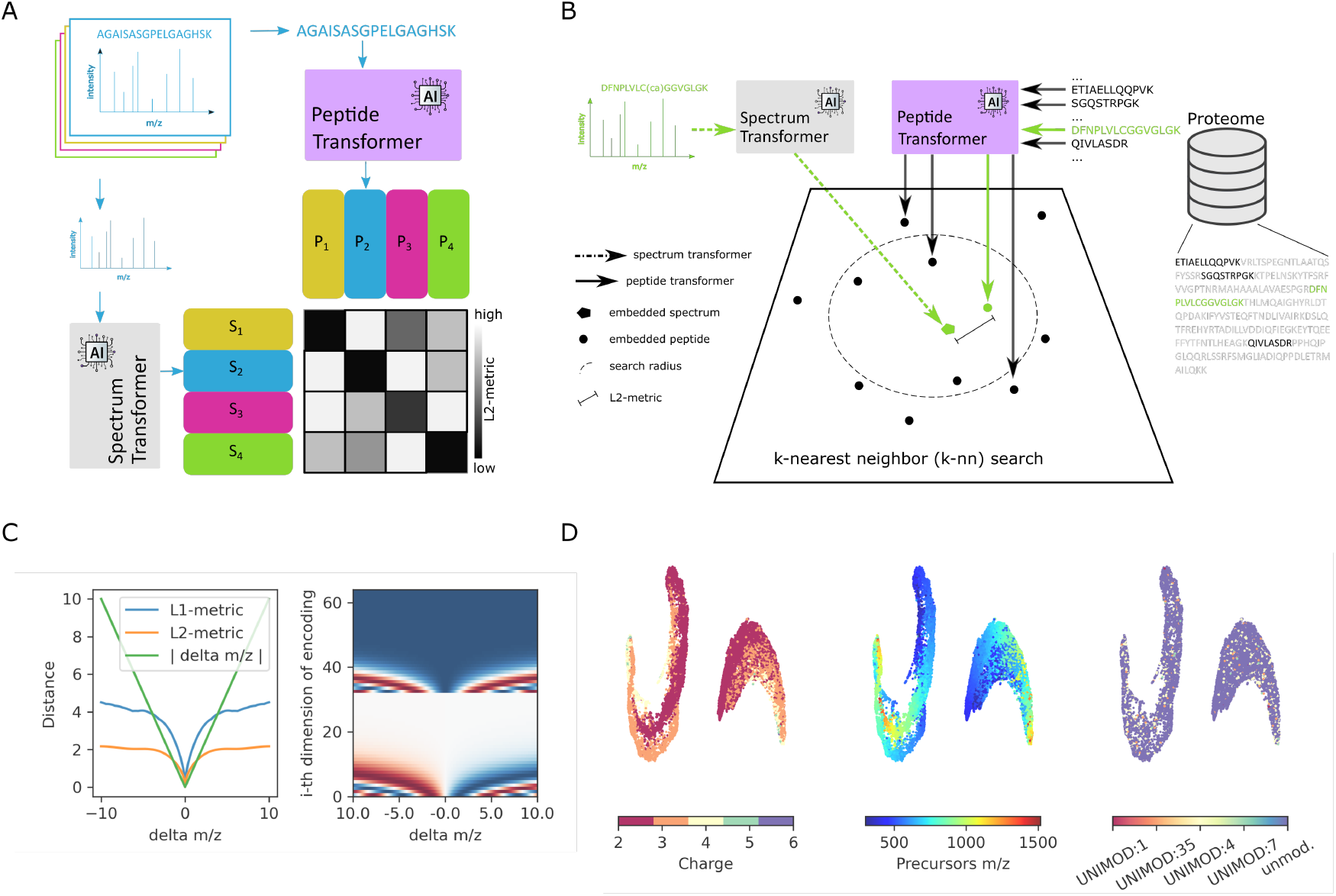
yHydra jointly embeds spectra and peptides. **A**: Illustration of the yHydra architecture: Spectrum Transformer for spectra (grey) and Peptide Transformer for peptide sequences (purple), each embed to their respective embeddings (yellow, blue, pink, green) for which pairwise Euclidean distances (L2-metric) serve as loss during training. The pairwise distance metrics allows to update model parameters such that the diagonal minimized and the off-diagonal is maximized. **B**: Illustration of the open search of yHydra, using a trained Spectrum Transformer and trained Peptide Transformer. Both Transformers embed batches of their respective inputs into the joint embedding space (center) in which a k-nearest neighbor search finds peptide candidates within an entire protein database. An individual spectrum and its correctly matching peptide is highlighted (green). **C**: Left, comparison of the behavior of distances between the wavelet encoding of the m/z location of a single peak with respect to deviations from this location (−10 m/z to 10 m/z). Right, visualization of the change for each of the i-th dimension of the wavelet encoding of a single peak depending on m/z deviations. **D**: UMAP visualization of the manifold of joint embeddings for identified peptide spectrum matches. The manifold appears as two clusters, spectra (left) and peptides (right). Each manifold is colored according to either charge, precursor mass. or post-translational modification. While charge and precursor mass show a clear structure in the UMAP, this is not the case for PTMs, supporting the hypotheses that the identification of modified peptides is feasible by exploring neighborhoods in embedding space.

As mentioned above, comparing spectra with one another typically requires peak matching. Instead, we use wavelet encoding (originally developed alongside the Transformer architecture) to encode the m/z location of each peak. This serves two purposes. First, it allows to present a spectrum to our Spectrum Transformer and second, it retains the ability to separate peaks in close proximity (Fig. 1C). To demonstrate this we offset two hypothetical peaks shown by the delta m/z on the x-axis (Fig. 1C). The change of each dimension of the 64 dimensional wavelet encoding is visualized (Fig. 1C, right panel) in comparison to applying L1– or L2-between the two peaks (Fig. 1C, left panel). This shows the information content and responsiveness of the peak encoding, giving the Spectrum Transformer its ability to input each peak and resolve its location.

Finally, because the learned joint embedding yields real-valued vectors of fixed size the entire toolbox of machine learning and statistical tools is open to be used in conjunction with our embeddings. For example, we used UMAP to visualize the manifold of the embeddings from identified PSMs (Fig. 1D). The two-dimensional manifold of both peptide and spectrum embeddings of identified PSMs is colored according to charge, precursor mass, and modifications. The embeddings are ordered from smaller to larger charges. They are also sorted by precursor masses. This indicates that representations retain information about those properties. In contrast, the PTMs are largely uniformly scattered, which makes the embedding suitable for an open search.

### yHydra enables ultra-fast open searching

MS-based proteomics is able to characterize proteoforms due to PTMs. However, accounting for PTMs increases the search space and thus an open search is needed to cope with the increased search complexity. Here, we implement an open search as a sub-task for our foundation model. Our open search allows for a delta mass that is a mass difference between the unmodified peptide from the reference proteome and any modified version of that same peptide. This delta mass can give rise to the underlying PTMs, specifically we observe oxidations (+16 Da) and carbamidomethylations (+57 Da) and combinations thereof (Fig. 2F). For the open search, yHydra starts by generating a set of tryptic peptides for a selected protein database (Fig. 1B). For each peptide, the trained Peptide Transformer infers a peptide embedding. Similarly, for each MS/MS spectrum in a run, the spectrum embedder infers a spectrum embedding. Subsequently, these spectrum embeddings are queried against the entire set of peptide embeddings using a k-nearest neighbor (k-NN) search. Note that all spectrum embeddings of an entire run (typically multiple ten-thousands) are queried simultaneously against the entire database in a single call to achieve lowest possible search times. To be able to search spectra against dedicated mass buckets (e.g. to select between a close, narrow or open search) while performing a single query per run we developed a multiplexed k-NN search (Methods). As a result, the yHydra search only takes seconds when using the GPU-acceleration, and thus being faster than MSFragger (Table 1). In this experiment we chose a k of 50 (k is a user-defined parameter, see Discussion) and only the 50 closest peptides are then scored by constructing a theoretical spectrum, matching peaks, and subsequent false discovery rate estimation using a target-decoy approach (Fig. 2E).

**Table 1:** Runtimes of yHydra and MSFragger on PXD007963 using a compute node with an A100 (NVIDIA) GPU and a EPYC (AMD) CPU.

|  | yHydra | MSFragger |
| --- | --- | --- |
| per run [seconds] | **113.3** | 183.0 |
| total runtime [seconds] | **340.0** | 549.0 |

**Figure 2:**
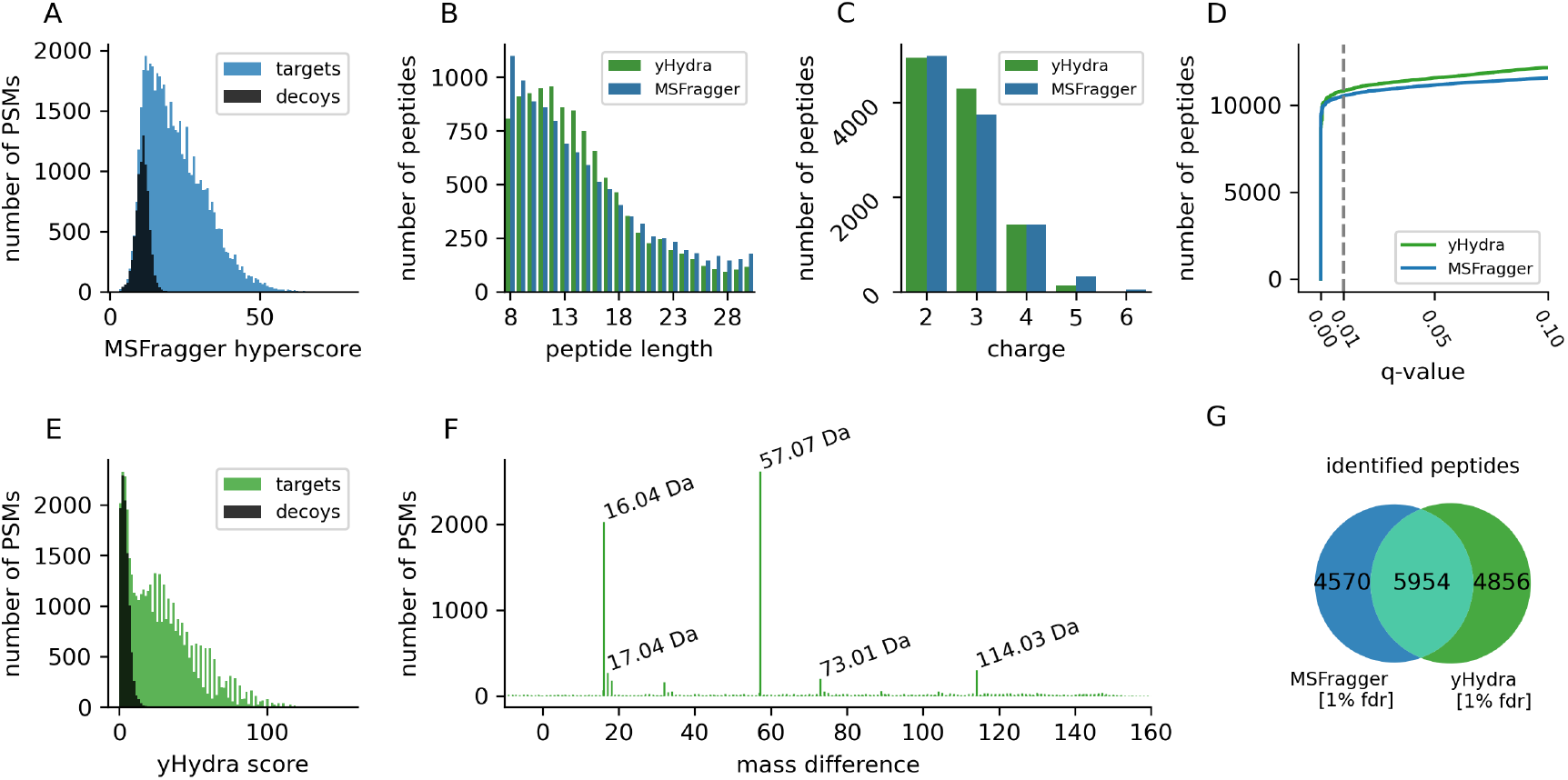
yHydra enables ultra-fast open searching, allowing an unrestrictive characterization of post-translational modifications in cyanobacteria. **A**: Target-decoy distribution of MSFragger hyperscores. **B**: Peptide lengths distribution according to MSFragger (blue) and yHydra (green) identifications. **C**: Precursor charge distribution according to MSFragger (blue) and yHydra (green) identifications. **D**: q-values estimate the FDR (x-axis) in comparison to number of identifications (y-axis) for yHydra (green) and MSFragger (blue) **E**: Target-decoy distribution of yHydra score. **F**: yHydra matches spectra to peptides in a wide mass range, the resulting delta masses between precursor mass and peptide mass are shown here. The five most common delta masses are labeled, including oxidation (Ox) +16 Da, cysteine carbamidomethylation (CAM) +57 Da, and their combinations, +73 Da is Ox+CAM and two CAMs is +114 Da. **G**: Venn diagram representing a unique set peptides identified by MSFragger (blue, 1% FDR), the intersection of identified peptides from both methods (turquoise; 1% FDR) and a unique set of peptides only identified by yHydra (green, 1% FDR).

Here, we search a sample from cyanobacteria (PXD007963) and compare open searches by yHydra and MSFragger (Fig. 2). Both methods identify peptides resulting in similar peptide length and charge distributions (Fig. 2B,C). Our yHydra score (Fig. 2E) resembles a bimodal distribution and thus separates targets and decoys better than MSFragger’s hyperscore (Fig. 2A). Due to the open search mode each PSM is assigned with a delta mass (Fig. 2F). Finally, yHydra identified 10,810 peptides at 1% FDR (Fig. 2D). In comparison, MSFragger identified 10,524 peptides at 1% FDR while both search engines share 5,954 of these peptides (Fig. 2G).

### yHydra enables error-tolerant search via gradient descent

As an additional sub-task for our foundation model we implemented an error-tolerant search by using a gradient descent in the learned embedding space of yHydra. An error-tolerant search identifies peptides that deviate from the given reference proteome. A proteome is context-matched with a finite collection of protein sequences. However, real protein sequences can have sequence variations. An error-tolerant search uses the reference proteome as guidance while being able to deviate from the contained protein sequences and thus accounts for mutations. The gradient descent inputs PSMs with non-zero delta mass and a minimum score, being an user-adjustable parameter. Delta masses (mass difference between precursor and the candidate peptide) come from the output of our open search, see previous section. In particular, each descent starts with a candidate peptide(Fig.3A, indicated step-0). Along the gradient descent, peptides are updated based on the gradient information (Fig.3A, green boxes) when gradient steps are made towards the MS/MS spectrum embedding (Fig.3A, blue box). Based the L2-gradient we update the peptide embedding and subsequently also the peptide sequence (Fig.3A). The search terminates if the delta mass is zero or maximum number of steps is reached. The search was successful if the updated peptide has an improved score (compared to the initial candidate peptide) (Fig.3A, dark green box).

To explore a wide range of peptide decodings by considering multiple alternative gradient descents (due to possible multiple local minima in embedding space, (Fig.3A)) we implemented a beam search using yHydra (Fig.3B). For the beam search, at each updating step and looking at the reward matrix (Fig.3B, purple matrix) we follow multiple potential AA-changes, so-called beams. In contrast to a simple greedy search where only the single best change per step is kept. Two beams are illustrated, the correct beam (Fig.3B, green arrows) and one alternative beam (Fig.3B, red arrows). Both beams are specific realizations of AA-changes (Fig.3B, indicated steps 0-3). All beams per PSM are kept and if the resulting peptide has a non-zero delta mass its score appears in the score-sorted list of beams (Fig.3B, list of beams). Finally, per PSM the top scoring peptide with zero delta mass is reported in the results of the error-tolerant search (Fig.3B, list of beams). The beam search explores multiple potential solutions (beams) and we report the overall best solution as the identified peptidoform (Fig.3B, top-scoring peptide in list of beams) per PSM.

**Figure 3:**
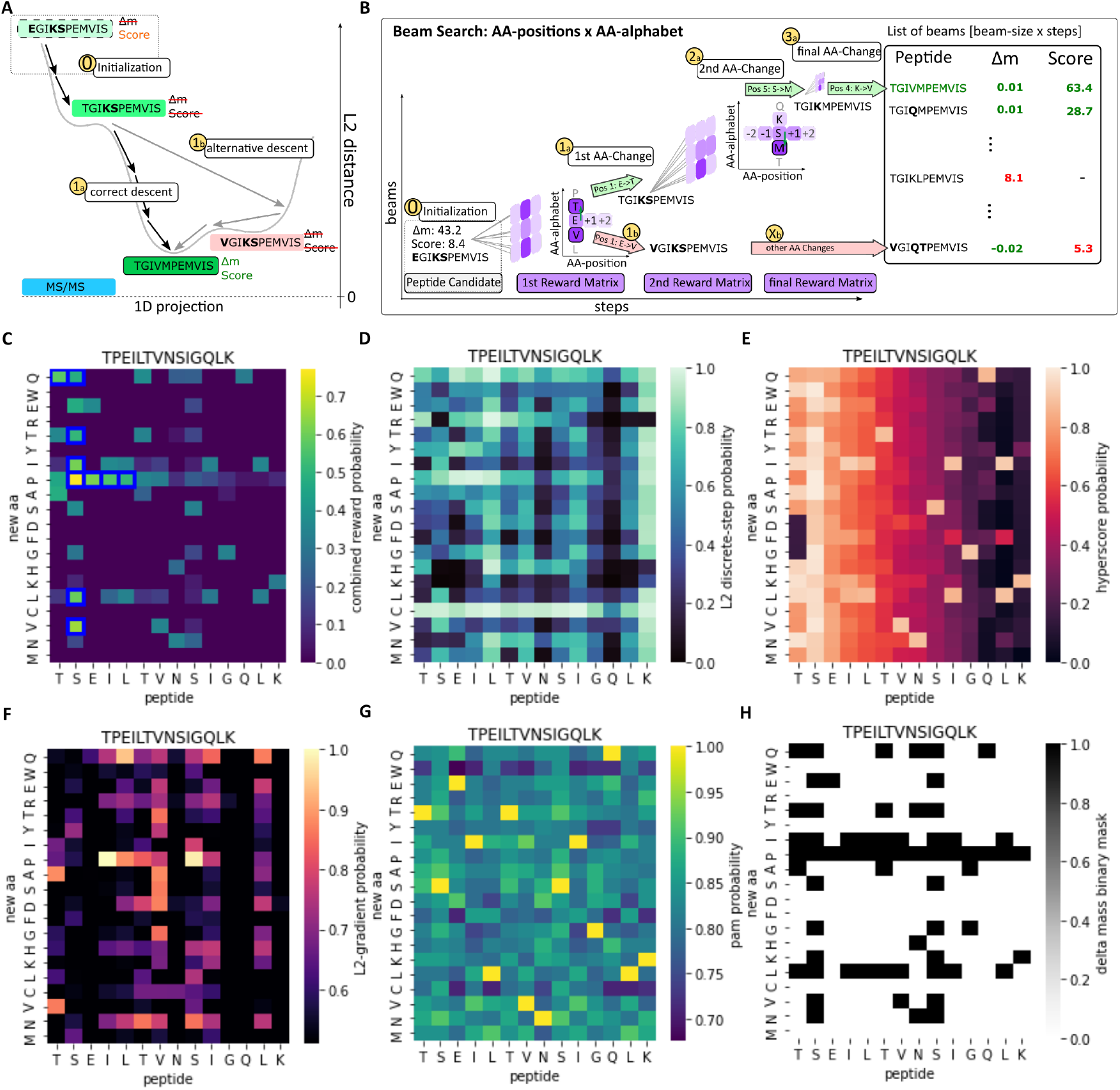
yHydra facilitates an interpretable error-tolerant search via gradient descent. **A**: Illustration of the gradient descent updating a candidate peptide EGIKSPEMVIS (0-step) towards the correct peptide TGIVMPEMVIS (dark green box) via AA-changes informed by the L2-gradient descent towards the MS/MS-embedding (blue box). A correct descent (1a-step) and alternative descent (1b-step) illustrate two alternative local minima. **B**: Overview of the beam search exploring two alternative beams: the correct beam (green arrows) and one alternative beam (red arrows). Beams are based on the reward matrix (purple) in each decoding step. The search initializes with a peptide candidate (0-step) and performs AA-changes (1a,2a,3a-steps) decoding the correct peptide TGIVMPEMVIS, ranked as top-scoring peptide in the list of beams. Exploration of an alternative beam (1b to Xb-steps) terminates with a low scoring peptide appearing at the bottom of the list of beams. **C**: Combined reward matrix, showing the probability of changing one AA of the peptide (x-axis) into any other AA from the AA-alphabet (y-axis), beams of beam-size 10 are indicated (blue boxes), where the correct change (S *→* P at position 2) has the highest probability. **D-G**: for interpretability, comparing the four AA-change reward matrices: L2-based-, hyperscore-based-, L2 gradient-based– and PAM-based probabilities that we are able to compute using yHydra. **H**: delta mass binary mask showing allowed AA-changes based on possible AA-changes restricted by the delta mass (between candidate peptide and precursor). For **C-H**, the true peptide is shown as panel title, whereas the current candidate peptide AAs are labels on the x-axis.

### yHydra is interpretable by looking at the AA-change reward matrices of the beam search

To interpret yHydra we take a closer look at the reward matrix of the gradient descent search (Fig.3B, purple matrix) described in the previous section. yHydra is comprised of two embedding blocks, the Spectrum Transformer and the Peptide Transformer (Fig.1A). Both are differentiable with respect to their inputs. This is essential to train both models but also provides means to interpret them. To interpret the peptide embedding we calculate the gradient of a given peptide embedding relative to its matching spectrum embedding (i.e. the partial derivative for each AA-position). Using the learned AA-encoding matrix we can further project this partial derivative onto a matrix that has the shape of the AA-alphabet times the peptide length (Fig.3F), see Methods.

As baselines we compare additional types of reward matrices that express AA-changes (Fig.3D-H). Therefore, we computed the Point Accepted Mutation (PAM) matrix (Fig.3G), which generally describes mutation probabilities of protein AA-changes, i.e. independent from our model. Also, we employ a binary delta mass matrix which constraints all possible AA-changes given a certain delta mass (Fig.3H). Lastly, we compute reward matrices based on the L2-distance and the hyperscore (Fig.3D+E). These two require exhaustively changing each AA at each position and store the changes of either L2 (Fig.3D) or the hyperscore (Fig.3E) as matrices. Note, these latter two matrices are more expensive to compute than our L2-gradient based matrix. For the beam search above we rely on the combined reward matrix (Fig.3C) which is a combination of the gradient matrices (Fig.3F-H), see Methods.

We can interpret yHydra by looking at the combined reward matrix (Fig.3C). This matrix is a visualization of how a current peptide, in this case TSEILTVNSIGQLK, should be changed. For this particular candidate peptide the single highest reward change is changing S (at position 2) into P, which results in the correct peptide TPEILTVNSIGQLK (Fig.3C).

### yHydra boosts peptide identifications for a monoclonal antibody via error-tolerant searching

Our gradient descent (gd) error-tolerant search can be used either as standalone or as post-processing to existing open search search engines. Here, we demonstrate both and compare i) yHydra open search with boosting via error-tolerant search via gradient descent ‘yHydra gd’ and ii) MSFragger open search and subsequent post-processing of MSFragger results via our gradient descent strategy, which we annotate as ‘MSFragger gd’ subsequently (Fig.4). As described above (Fig.3A) the error-tolerant search is initialized from candidate PSMs that have a substantial delta mass (Fig.3A) and a minimum score (Fig.3B) both are parameters of the error-tolerant search, see Methods. The gd error-tolerant search is formulated as an optimization algorithm that aims to reduce the absolute delta mass while simultaneously maximizing the score for the PSMs with their updated peptides. The gd error-tolerant search is formulated as an optimization algorithm that aims to maximize the score for the PSMs while removing the delta mass (i.e. delta mass being zero) by updating the peptide sequence based on the gradient information. This –explaining-away of delta mass for a better score– can be seen (orange dashed box in Fig.4A), while most PSMs before the gd search are outside of orange box they enrich after the gd search (MS-Fragger gd and yHydra gd) in that box. Simultaneously these same PSMs receive a better scoring, shifting the score distribution of initial candidate PSMs to higher scores for both gradient descent searches MSFragger gd and yHydra gd (Fig.4B). Here, we searched a tryptic digest of a monoclonal antibody (from MSV000079801) against heavy and light chains of human immunoglobulin sequences that are publicly available in UniProt [31]. To control for the FDR we kept decoy PSMs (Fig.4A-B) throughout the workflow and evaluated q-values (Fig.4C) based on these decoys that underwent gradient descent search similar to true peptides. Actual hits are peptides contained in the complete assembly of the monoclonal antibody MSV000079801 [28] serving as ground truth here. As a result, the peptide identifications from the monoclonal antibody are boosted, from 39 to 43 peptides for MSFragger to MSFragger gd and from 50 to 55 peptides for yHydra to yHydra gd (Fig.3C+D), each at 5% peptide FDR.

**Figure 4:**
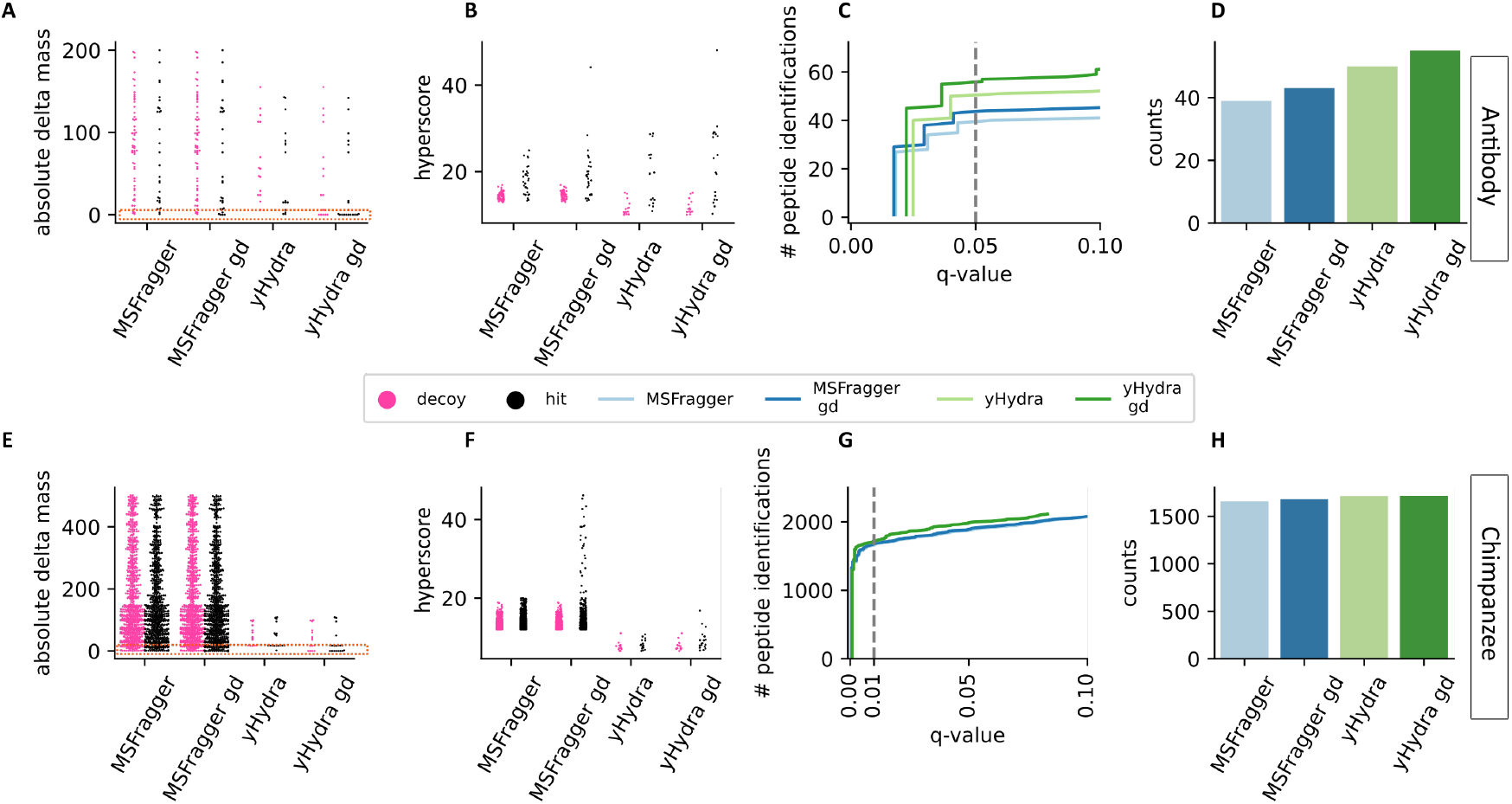
yHydra enables error-tolerant searching of a monoclonal antibody and chimpanzee plasma. **A-D**: Comparisons of the antibody searches by MSFragger and yHydra with additional gradient descent mode (gd). **A**: Absolute delta masses show a better enrichment (orange box) of hits (black) at around zero compared to decoys (pink), comparing MSFragger, yHydra with and without gradient descent (gd). **B**: hyperscores of gd-candidates before and after gradient descent (gd). **C**: q-value curve showing the number of peptide identifications for MSFragger and yHydra before and after gradient descent (gd). Dashed line indicates the number of peptide identifications at 5%FDR peptide level (q-value), shown in **D. E-H**: same as **A-D** but for the chimpanzee plasma sample.

### yHydra error-tolerant search identifies chimpanzee peptides in a cross-species search

Here, we searched a tryptic digest of a chimpanzee blood serum sample against the human proteome in an error-tolerant manner (Fig.4E-H). Gentic differences between chimpanzee and human are well understood on the genome level, but experimental evidence for the majority of the predicted chimpanzee proteins is missing [30]. In a first pass, we performed open searches using yHydra or MSFragger and the resulting PSMs served as candidates for the gradient descent strategy. These second passes are called yHydra gd and MSFragger gd respectively (Fig.4E+F). As ground truth we used predicted protein sequences from genome translation of protein-coding genes of the chimpanzee genome [30, 29]. Specifically, an identified peptide is counted as hit when it is contained in the aforementioned predicted chimpanzee proteome. Similarly, we keep decoy hits from the reversed human protein sequences (open search candidates) throughout the gradient descent workflow and they are counted as decoy hits if they appear in the reversed chimpanzee protein sequences. This allows us to control for the FDR (q-value) (Fig.4G+H).

As a result, MSFragger identified 1658 peptides and yHydra found 1715 peptides (Fig.4H). For the error-tolerant searches, MSFragger gd finds 1678 peptides while yHydra gd yielded 1717 peptides at 1% FDR.

## Discussion

We propose a new approach of how to embed peptides and spectra of MS-based proteomics jointly and thus provide a foundation model to solve various sub-tasks. Using our model, we implemented an open search to unrestrictedly characterize post-translational modified peptides. Besides, we implemented an error-tolerant mode, that we evaluated as a standalone but also as post-processing to an existing algorithmic search engine. Our foundation model serves as a platform to implement further sub-task either by implementation or fine-tuning. In particular, we provide trained model weights for the two instruments Q Exactive and timsTOF, as well as for tryptic and non-tryptic peptides (Supplementary Material). We evaluated our model on relevant proteomics datasets, identifying proteoforms in a monoclonal antibody digest and a chimpanzee serum sample.

Training of yHydra makes use of historical data, consisting of previously identified peptide spectrum matches (PSMs). We directly used PSMs for the contrastive loss function because each mini-batch consists a set of matching PSMs (diagonal of L2-matrix in Fig. 1A) and non-matching PSMs as negative training examples. We found that a random set of PSMs per mini-batch is sufficient. Additional MS1-precursor matching per mini-batch is not improving the model performance any further. For the open search we use a GPU-accelerated k-NN search (Fig. 1C). The number of neighbours k is a parameter, configurable by the user. Generally, a smaller k tends improves runtime, while a larger k tends to improve sensitivity. However, as a heuristic and depending on the dataset, a larger k may also result in low-scoring PSMs in the final target-decoy scoring, which effectively may reduce the number of identified peptides. Also, for an open search a larger k (larger than 100) is advisable since not only peptides but also their peptide species with various delta masses need to be considered.

To present spectra to the Spectrum Transformer we use wavelet encoding (Fig. 1C). We could show, that the wavelet encoding is able to resolve two peaks in close proximity. In particular, we show that the L2 distance applied directly on a peak encoding (before the Transformer) is approximating the actual distance between the two peaks and is highly responsive especially at closer distances for better resolution (Fig. 1C). An open search by yHydra is possible because our embeddings are largely indifferent across PTMs, i.e. a peptide sequence can be identified regardless of specific PTMs (however the reported delta masses can give rise to PTMs). Specifically, this can be seen from the respective UMAP visualizations (Fig. 1D) and in the search results themselves and ultimately from the resulting delta mass profile of identified peptides (Fig. 2A).

For the open search we evaluated yHydra against MSFragger, searching a sample of tryptic peptides from cyanobacteria. Despite the two methods employ different approaches, both show comparable results with more identified peptides in case of yHydra. MSFragger is an algorithmic approach based on a fragment-index, whereas ours is making use of the joint embeddings in Euclidean space to identify k peptide candidates in a k-NN search. These k candidates are subsequently scored (Fig. 2E). When comparing the two score distributions of yHydra (Fig. 2E) and MSFragger (Fig. 2A) we see that yHydra has a better separation between target and decoy PSMs. Using yHydra for open searching additionally results in PSMs with delta masses. Each delta mass quantifies the difference between the MS1 precursor mass and the identified peptide mass. This delta mass is the aggregated mass of potential PTMs or due to sequence variants. A summary of all delta masses in the cyanobacteria sample is summarized as a delta mass profile (Fig. 2F). Here, we observe PTMs like an oxidation (Ox) +16 Da, cysteine carbamidomethylation (CAM) +57 Da, or combinations thereof. For example, 73 Da is Ox+CAM and two CAMs result in +114 Da (Fig. 2F).

We implemented an error-tolerant search using our foundation model yHydra. Using our beam search we explored alternative AA-changes (Fig.3A+B) due to the 2nd-, 3rd– and xth-best value in the combined reward matrix (Fig.3C) for a given peptide candidate. Taking into account these alternative updates (i.e. alternative beams) increases our chances to decode the correct peptide (Fig.3A, correct descent 1b). The final peptide is identified by scoring each decoded peptide with zero delta mass (Fig.3B). The number of alternative beams is a sensitivity-to-runtime trade-off, selectable by the user. For the two error-tolerant experiments (monoclonal antibody and chimpanzee plasma) we were able to decode peptides that score higher than their database-derived counterparts (Fig.4B+F). While these decoded peptides also result in an enrichment of zero delta masses after they have been decoded (Fig.4A+E, orange box). The increase of scores and vanishing of delta masses is predominantly true for hits against the ground truth sequences while less pronounced for decoy peptides. Ultimately, this suggests that our error-tolerant search using a gradient descent is a capable strategy to explore peptidoforms in such challenging datasets.

Altogether, our foundation model provides a powerful framework for MS-based proteomics. In our experiments for the various sub-tasks it surpasses other algorithmic approaches and results in robust peptide identification. We foresee, that our foundation model may also be extended to additional sub-tasks, as it is not only limited to the cases shown in this work. For example, *de novo* sequencing may be possible within the joint embedding space, as it would be related to our gradient descent approach and as dedicated Transformer models exist for *de novo* sequencing [21]. Further sub-tasks may be multi-omics integration by employing the embeddings and passing them to a downstream machine learning model, similar to the UMAP of our embeddings. Lastly, an additionally trained gene sequence Transformer could be a promising approach for tackling the six-frame translation problem in proteogenomics [14].

## Methods

### Preprocessing and encoding of spectra and peptides

The intensities are normalized such that the intensities (*I*_1_, …, *I*_*l*_, …, *I*_*N*_) are a unity-length vector (i.e. scaled by 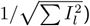). For each peaks list, up to 500 peaks are kept. Otherwise, if the list contains more than the maximum number of peaks, we keep peaks according to the top-500 highest intensities or append tuples of (0.0,0.0) if the peaks list contained less peaks.

Both spectra and peptides are fed to Transformer architectures, hence we adapt the original positional encoding (also called wavelet encoding) and extend it to suit the context of spectra and peptides.

For the positional encoding of the peaks list of the peaks list (*mz*_1_, …, *mz*_*l*_, …, *mz*_500_), each *mz*_*l*_ is multiplied by the radiant rate *r*_*i*_ of the *i*-th dimension of the encoding vector *s*: *s*_*li*_ = *mz*_*l*_ *·r*_*i*_, where *r*_*i*_: = 10000.0^*−i/d*^. Furthermore, 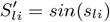 if *i* is an even integer and 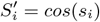 if *i* is odd. For the final spectrum encoding *S*, each positional encoded *mz*_*l*_ is scaled by its respective intensity: 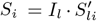. Hence *S* is a tensor of size (number of peaks, *d*) = (500,64).

For each peptide sequence, we enumerate each amino acid (starting from zero, up to peptide length n – 1) these integer positions *p*_*n*_ are then multiplied by a radiant rate *r*_*i*_ (same definition as above) of the *i*-th dimension of the peptide positional encoding vector *p*: *p*_*li*_ = *p*_*l*_ *· r*_*i*_. Up to n=42 amino acids are allowed, if the peptide is shorter it is padded with zeros. Hence, the peptide positional encoding vector has a size of (42,64). Furthermore, the amino acids are replaced by unique indices (starting from 0 up to 25, with zero being a padding character) and function as the index into a lookup table of size (alphabet size, d) = (26, 64), where the parameters of the lookup table are trainable and trained together with the model. For the final peptide sequence encoding *P*, each positional encoded *p*_*i*_ is multiplied by its amino acids embedding (from the lookup table): *P*_*li*_ = *p*_*li*_ *· aa*_*i*_. Hence *P* is a tensor of size (maximum peptide length, d) = (42,64).

### Training of yHydra

#### Training data

The Transformer models of yHydra were trained on 19,991,263 PSMs (retrieved as USIs) from the following 67 repositories: PXD000702, PXD001072, PXD001344, PXD001351, PXD002054, PXD002147, PXD003094, PXD003261, PXD003364, PXD003556, PXD003718, PXD003779, PXD003916, PXD003976, PXD004398, PXD004825, PXD005009, PXD005117, PXD005196, PXD005306, PXD005341, PXD005654, PXD005744, PXD006033, PXD006084, PXD006316, PXD006375, PXD006389, PXD006645, PXD006823, PXD006836, PXD008592, PXD008602, PXD008622, PXD008647, PXD008667, PXD008895, PXD009387, PXD009665, PXD009698, PXD009713, PXD010000, PXD010641, PXD010827, PXD011042, PXD011583, PXD011712, PXD011714, PXD011984, PXD012827, PXD013274, PXD013304, PXD013684, PXD013711, PXD013712, PXD013724, PXD013890, PXD013897, PXD015153, PXD015296, PXD015698, PXD016833, PXD016846, PXD017308, PXD018714, PXD019095, PXD019134. We specifically selected repositories of non-model organisms (excluding the top-10 most commonly studied organisms in terms of counts of repositories) to reduce the bias towards certain proteome-specific sequence patterns. Furthermore, we only included data acquiered on Q Exactive.

#### Pairwise contrastive loss of yHydra

The loss of yHydra is inspired by the recent approach of CLIP [26] and is based on the idea to directly calculate a contrastive loss based on the pairwise distances within each mini-batch (table 2 and Fig.1A). We found this approach has major advantages over previous types of contrastive-losses or triplet-losses while virtually having none of their shortcomings. Most importantly the pairwise contrastive loss does not not require the creation of artificial negatives as they naturally occur due to the mixed pairs of distances (i.e. off-diagonal elements in Fig.1A). This is not only more elegant than previous contrastive loss formulations but also makes the network learn at anytime, whereas for older types of contrastive losses typically hard-negative mining was essential to get decent training results. Furthermore, we extended this idea by adding label smoothing which should allow the model to also learn from the specific but small distances that mixed negatives naturally have. Label smoothing allows the model to also learn from this regime (i.e. off-diagonal elements in Fig.1A).

**Table 2:**
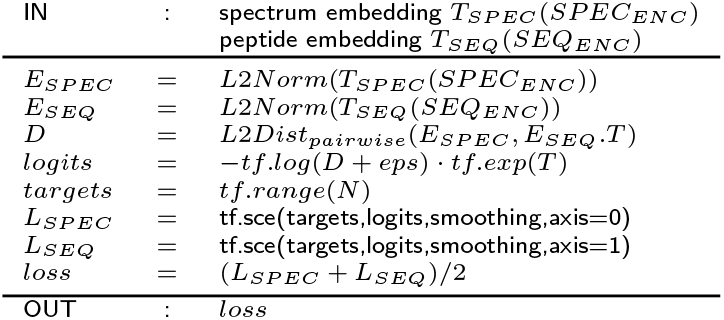
Tensorflow-/Numpy-like pseudocode of the calculation of the pairwise loss between spectra and peptides. The Transformer models *T*_*SPEC*_ and *T*_*SEQ*_ yield embeddings *E*_*SPEC*_ and *E*_*SEQ*_ each of size (batch-size=64, embeddings-size=64). The spectrum encoding *SPEC*_*ENC*_ and peptide sequence *SEQ*_*ENC*_ are described in the main text. Parameters are set to eps=0.001, smoothing=0.1 and T=3.0 and tf.sce is the sparse cross entropy between targets and logits according to the the distance matrix D (illustrated in Fig. 1A).

### yHydra algorithm

#### Multiplexed k-NN search of mass buckets for closed, narrow and open searches

Our multiplexed k-NN search allows us to search all spectra embeddings against all peptide embeddings within the same search call while at the same time only spectra against theoretically possible peptides (e.g. determined by certain combinations of precursor mass and respective peptide masses) are searched. Therefore we divide the peptides in the database into buckets according to their theoretical mass. Hence, for a closed search we could have a thousand of small buckets (of +/-1 Da width) and, in contrast, for the open search we have a few but wide buckets (e.g. +/-500 Da width). Each bucket gets a unique vector assigned, which is appended to the peptide embeddings in that bucket (i.e. similar to an ‘address’ vector). Subsequently, the query embeddings, which are supposed to be searched against a specific bucket also gets the respective address vector appended. Effectively, the L2-norm between the embeddings is dictated by their common ‘address’-vector because only those with a common address-vector have meaningful intra-buckets L2-distances but comparatively high inter-bucket L2-distances. Ultimately, this allows us to achieve multiple mass-compliant search calls while really only performing a single search.

#### GPU-accelerated peak matching and PSM scoring

The core algorithms of yHydra are GPU-accelerated (i.e. neural networks and k-NN search by faiss [27]). To further speed up the runtime of yHydra we developed a GPU-accelerated peak-matching and PSM scoring (table 3). The idea is to simultaneously score a batch of 64 spectra against their respective k-candidates, i.e. k=50, which means for 3,200 PSMs peaks are matched and scored in parallel.

**Table 3:**
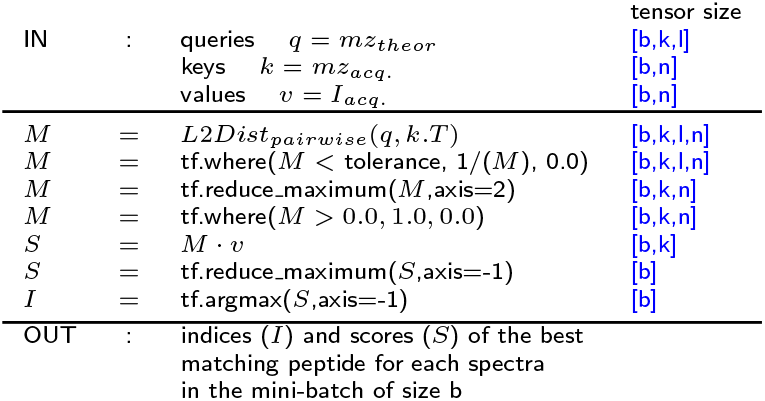
Tensorflow-/Numpy-like pseudocode of peak matching and scoring for PSMs. The inputs are a mini-batch of b spectra, with n peaks considering their mz-values *mz*_*acq*._ and intensities *I*_*acq*._. Furthermore, list of k candidate peptide (result of the k-NN search) is considered as theoretical ions with up to *l* mz-locations *mz*_*theor*_, see Methods for details on parameters.

#### yHydra error-tolerant search via gradient descent

To update a candidate peptide we used the gradient in the Euclidean space. The gradient is the partial derivative of the difference between the peptide embedding and the spectrum embedding: 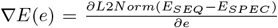, along each dimension *e* of the embedding space. By multiplying that gradient with the pairwise difference matrix of learned aa-encodings, which has a shape of (alphabet-size,alphabet-size,embedding-size) we gain the final reward matrix of gradient-based aa-changes: 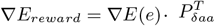. The reward matrix *∇E*_*reward*_ has the shape of maximum peptide length times alphabet-size. A particular reward matrix evaluated for the peptide TSEILTVNSIGQLK is displayed in the main text.

#### Beam search

The beam search helped us to explore many possible updates by also accounting for secondary alternative gradient directions. Each beam starts from the L2-gradient based reward matrix *∇E*_*reward*_ as it provides alphabet-size times peptide length potential updates in a single step. This reward matrix can be seen as a ranking of updates. We set the beam size as percentage of the size of the reward matrix. For example, a peptide of length 14 and considering the alphabet-size 20, using beam size of 10% results in 28 updates for the next step. We tested different beam sizes and selected a beam size of 20% with three update steps for our experiments.

### Open search parameters for yHydra and MSFragger

For both search open engines (yHydra and MSFragger) the raw files qe2 03132014 1WT-1, qe2 03132014 5WT-2, and qe2 03132014 13WT-3 were searched against the Syn-PCC7002 Cbase.fasta by considering tryptic peptides with up to 1 miscleavage of length between 7 and 42 amino acids. In both searches a minimum delta mass of –150.0 Da and a maximum delta mass of 500.0 Da is considered.

For yHydra, the matching tolerance was set to 0.01 Da and a mimimum of 4 matching peaks for a PSM was required. For scoring, the globally highest top-100 peaks per spectrum are considered. For each pepetide, b- and y-ions are calculated and up to 200 fragment ions are considered (any excess ions are discarded starting from higher charge states). For MSFragger, we used the standard search parameters of MSFragger version 3.3, the maximum peptide length of 42 and allowing up to 1 miscleavage. For both methods, the PSM-level FDR was set to 1%.

## Data availability

Proteomic data were downloaded from public repositories PXD000702, PXD001072, PXD001344, PXD001351, PXD002054, PXD002147, PXD003094, PXD003261, PXD003364, PXD003556, PXD003718, PXD003779, PXD003916, PXD003976, PXD004398, PXD004825, PXD005009, PXD005117, PXD005196, PXD005306, PXD005341, PXD005654, PXD005744, PXD006033, PXD006084, PXD006316, PXD006375, PXD006389, PXD006645, PXD006823, PXD006836, PXD008592, PXD008602, PXD008622, PXD008647, PXD008667, PXD008895, PXD009387, PXD009665, PXD009698, PXD009713, PXD010000, PXD010641, PXD010827, PXD011042, PXD011583, PXD011712, PXD011714, PXD011984, PXD012827, PXD013274, PXD013304, PXD013684, PXD013711, PXD013712, PXD013724, PXD013890, PXD013897, PXD015153, PXD015296, PXD015698, PXD016833, PXD016846, PXD017308, PXD018714, PXD019095, PXD019134 to train yHydra. Furthermore, public data from PXD007963 was downloaded to evaluate yHydra. For the monoclonal antibody benchmark, data was downloaded from MSV000079801 via MassiveKB. The Chimpanzee plasma data is available as a Zenodo dataset [32].

## Code availability

An open source implementation with command-line instructions is publicly available (under MIT license) at https://gitlab.com/dacs-hpi/yHydra. A separate open source repository for training yHydra is available at https://gitlab.com/dacs-hpi/yHydra_train.

## Acknowledgements

This work is supported by a European Research Council (ERC) grant (eXplAInProt, 101124385) to BYR.

## Competing interests

TA is employed by Johnson & Johnson Innovative Medicine.

## Notes

### Summary of Updates

Added Figures 3+4 and corresponding sections in the main text.

